# The Mycobacterium lipid transporter MmpL3 is dimeric in detergent solution, SMALPs and reconstituted nanodiscs

**DOI:** 10.1101/2024.05.06.592688

**Authors:** Sara Cioccolo, Joseph D. Barritt, Naomi Pollock, Zoe Hall, Julia Babuta, Pooja Sridhar, Alicia Just, Nina Morgner, Tim Dafforn, Ian Gould, Bernadette Byrne

**Affiliations:** Department of Life Sciences, Imperial College London, Exhibition Road, South Kensington, London, SW7 2AZ, UK; Department of Chemistry, Molecular Sciences Research Hub, Imperial College London, Shepherd’s Bush, London W12 0BZ, UK; School of Biosciences, University of Birmingham, UK; Division of Systems Medicine, Imperial College London, London, UK; Institute of Physical and Theoretical Chemistry, J.W. Goethe-University, Frankfurt am Main, Germany

**Keywords:** MmpL3, resistance, nodulation and cell division protein, transporter, mycolic acid, oligomeric status

## Abstract

The mycobacterial membrane protein large 3 (MmpL3) transports key precursor lipids to the outer membrane of Mycobacterium species. Multiple structures of MmpL3 from both *M. tuberculosis* and *M. smegmatis* in various conformational states indicate that the protein is both structurally and functionally monomeric. However, most other resistance, nodulation and cell division (RND) transporters structurally characterised to date are either dimeric or trimeric. Here we present an in depth biophysical and computational analysis revealing that MmpL3 from *M. smegmatis* exists as a dimer in a variety of membrane mimetic systems (SMALPs, detergent-based solution and nanodiscs). Sucrose gradient separation of MmpL3 populations from *M. smegmatis,* reconstituted into nanodiscs, identified monomeric and dimeric populations of the protein using laser induced liquid bead ion desorption (LILBID), a native mass spectrometry technique. Preliminary cryo-EM analysis confirmed that MmpL3 forms physiological dimers. Untargeted lipidomics experiments on membrane protein co-purified lipids revealed PE and PG lipid classes were predominant. Molecular dynamics simulations, in the presence of physiologically-relevant lipid compositions revealed the likely dimer interface.

## Introduction

Mycolic acids (MAs) are long, branched-chain fatty acids and a key component of the outer membrane of mycobacterial species including *Mycobacterium tuberculosis*, the causative agent of tuberculosis^1,2^. MAs affect the biophysical properties of the outer cell membrane, e.g. permeability. They are required for virulence^3^, and influence the host immune responses during mycobacterium infection^4^. The MAs are transported across the inner membrane in the form of trehalose monomycolate (TMM) which is then either used to form part of the trehalose dimycolate (TDM or cord factor) or covalently linked to the arabinogalactan-peptidoglycan layer as mycolyl arabinogalactan peptidoglycan (mAGP). The essential nature of the MAs has made them attractive drug targets, with several anti-tubercular agents targeting the MA biosynthesis pathways, including isoniazid and ethionamide ^5^.

More recently attention has focused on targeting the transport process whereby the MAs are transferred from the inner membrane, where they are made, to their final cellular destination in the outer membrane^6^. Mycobacterium membrane protein large 3 (MmpL3) has been identified as the key *M. tuberculosis* MA exporter protein^6,7^. MmpL3 is a member of the resistance, nodulation and cell division (RND) superfamily of ion dependent transporters and has been shown to move TMM across the inner membrane, an activity which can be blocked by small molecule inhibitors ^8^.

There are several structures available of MmpL3 both from *Mycobacterium tuberculosis* and the related bacterium *Mycobacterium smegmatis (Msmeg)* ^9–12^. These revealed that MmpL3 contains 12 transmembrane (TM) domains organised into two 6 helix bundles. MmpL3 also contains a large, structured (α-β-α-β-α-β) periplasmic domain (PD) made up of two sub-domains formed from extended loops that connect TM domains 1 and 2 and TM domains 7 and 8. There is close interaction between the two sub-domains with the first α-helix of each forming part of the associated sub-domain^9–12^. Crystal structures of MmpL3 from *M. smegmatis* in complex with a range of TB drug candidates revealed an inhibitor binding site within the TM domains, between the two helical bundles^9^. Additional crystal and cryo-EM structures of MmpL3 from *M. smegmatis* have been obtained in complex with the detergent molecule 6-DDTre, a homologue of TMM^9^, the native substrate TMM^11^, n-dodecyl-β-D-maltopyranoside (DDM)^13^, and a molecule of phosphatidylethanolamine (PE)^13^, as well as a structure of MmpL3 from *M. tuberculosis* in complex with a molecule of the detergent lauryl maltose neopentyl glycol (LMNG) ^12^. Taken together these structures reveal a large and flexible substrate binding site, predicted to be accessible from the outer leaflet of the membrane and which allows substrate transport via both the TM region and the PD. A combination of high-resolution structures in the apo and inhibitor bound states and molecular dynamics simulations suggests that a coordinated set of movements involving the TM domains and the PD mediates both proton translocation and substrate export ^14^.

Intriguingly, all of the structures of MmpL3 obtained to date indicate that MmpL3 is structurally and functionally a monomer. This is in contrast to most other RND transporters, the best characterised example of which is the bacterial multidrug efflux protein AcrB, shown to exist as a trimer both alone,^15^ and in complex with the other components of the tripartite efflux complex, AcrAB-TolC ^16^. The AcrB trimer is key for function of the protein ^17,18^, as export is coordinated via the alternating conformations adopted by the individual protomers, and is also likely important for association with AcrA and TolC to form the complete tripartite efflux complex. A trimer has also been reported for the related aminoglycoside exporter, AcrD^19^. In contrast, the hopanoid transporter HpnN is dimeric^20^. HpnN is responsible for movement of pentacyclic triterpenoid lipids that are structural and functional homologues of cholesterol, from the plasma membrane to the outer membrane of some gram-negative bacteria. However, the human Nieman-Pick C1 protein which plays a crucial role in cholesterol homeostasis appears to be monomeric^21^.

Here we used a variety of biophysical methods including analytical ultracentrifugation, native mass spectrometry and cryo electron microscopy, in combination with MD simulations and lipidomics analysis to explore the oligomeric status of MmpL3 from *M.smegmatis* and *M. tuberculosis.* Our data reveal that the protein is present as dimers in a range of different membrane mimetic systems, although monomer is always present too, indicating that the interactions supporting the dimer formation are relatively weak. Lipids may be key in stabilising the dimer.

## Materials and Methods

### Cloning and expression of the MmpL3 constructs

Plasmid pET22/42 encoding the C-terminal His_8_-tagged full length M*smeg* MmpL3 was obtained from Dr Zhujun Xu, from Professor Shu-Sin Chng’s group (National University of Singapore). Full length *Mtb* MmpL3 in pMA-T vector was kindly provided by Dr Oliver Adams, from Professor Simon Newstead’s group (University of Oxford). From the full-length constructs, C-terminally truncated versions were generated, by PCR amplification of aa 1-773 for ΔC Ms-MmpL3 and aa 1-753 for ΔC Mtb-MmpL3 and the products cloned into the same expression vector using NdeI and XhoI restriction enzymes. The primer sequences used are shown in Expanded View Table 1. The full-length and C-terminally truncated constructs were initially expressed in the BL21DE3 *ΔacrABEF* strain, generated and generously provided by Prof. Yamanaka (Hiroshima International University). For large scale expression, cells were grown in LB broth under selection at 37°C until the OD_600_ reached 0.5-0.7. Isopropyl β-d-1-thiogalactopyranoside (IPTG) was added to a final concentration of 1 mM and the cells were harvested by centrifugation after 4 hours, snap frozen and stored at –80°C until further use.

### Membrane preparation and protein purification in DDM or as SMALPs

Cell pellets were resuspended in Buffer A, 20 mM Tris HCl, 150 mM NaCl, 100 μg/mL lysozyme, 50 μg/mL DNaseI (DN25, Sigma-Aldrich) and EDTA-free protease inhibitor (SIGMAFAST^TM^, Sigma-Aldrich). Cells were lysed and the lysate was centrifuged at 12,000 g for 10 minutes at 4°C to remove unbroken cells and the supernatant then ultracentrifuged for 2 h at 200,000 g. Membranes were homogenised in Buffer B (20 mM Tris HCl, 150 mM NaCl, 10% v/v glycerol and protease inhibitor) and stored at −80°C. Membranes were solubilised by constant stirring for 1.5 hours at 4°C in DDM (Anatrace) at a final 1% w/v concentration. The extracted membrane proteins were then separated from non-solubilised material by ultracentrifugation at 200,000 g for 1h at 4°C. Ni^2+^-NTA Superflow resin (30410, Qiagen) was pre-equilibrated with Affinity Buffer (20 mM Tris HCl, pH 8.0, 150 mM NaCl, 10% v/v glycerol, 10 mM imidazole and 0.03% w/v DDM) and mixed with the membranes for 2 hours at 4°C. The resin mixture was then loaded onto a poly-prep/glass econo-column chromatography column (Bio-Rad) and allowed to drain by gravity. The resin was then washed with 10 column volumes (CVs) of Wash Buffer (20 mM Tris HCl, pH 8.0, 150 mM NaCl, 10% v/v glycerol and 0.03% w/v DDM), containing 30 mM imidazole. MmpL3 was then eluted from the column by addition of 5 CVs of Elution Buffer (20 mM Tris HCl, pH 8.0, 150 mM NaCl, 10% v/v glycerol and 0.03% w/v DDM and 300 mM imidazole) and the eluted protein was concentrated to ∼500 μL using 100 kDa MWCO centrifugal filters (Millipore Amicon^TM^ Ultra, Merck). The concentrated sample was then loaded onto a Superdex 200 10/300 gel filtration column (Cytiva) pre-equilibrated with SEC Buffer A (20 mM Tris HCl pH 8.0, 150 mM NaCl, 10% glycerol, 0.03% DDM). Fractions corresponding to main protein peak were then concentrated using 100 kDa MWCO centrifugal filters and either used fresh or stored at −80°C for future experiments.

For preparation of SMALPs, membranes obtained as described above were resuspended in SEC buffer B (20 mM Tris HCl pH 8.0, 150 mM NaCl) and diluted to a concentration of 60 mg/mL (wet weight of membranes). Membranes were then solubilised with gentle shaking for 2 hours at room temperature in a buffer containing 20 mM Tris HCl pH 8.0, 150 mM NaCl and 2.5% w/v SMA2000P. After centrifugation at 200,000 g for 1 hour at 4°C, the supernatant containing the soluble fraction was incubated with Ni^2+^-NTA resin overnight at 4°C, with rocking. The resin sample was then loaded into a poly prep/glass econo-column chromatography column (Bio-Rad) and the flow through allowed to drain out by gravity. The resin was initially washed with 20 CVs of imidazole-free SEC buffer B, followed by 10 CVs of SEC buffer B containing 10 mM imidazole and finally the SMALP-MmpL3 was eluted by washing the resin with 5 CVs of SEC buffer B containing 500 mM imidazole. The eluted sample was concentrated using 100 kDa MWCO centrifugal filters to a final volume of ∼500 μL prior to loading onto a Superdex 200 Increase 10/300 gel filtration column pre-equilibrated with SEC buffer B. Fractions from the peak were concentrated using 100 kDa MWCO centrifugal filters and either used fresh or stored at −80°C for future experiments.

### Reconstitution of the purified proteins into nanodiscs

The concentrated sample of detergent solubilised MmpL3 after elution from Ni^2+^-NTA column was mixed with Membrane scaffold protein 1 E3D1 (MSP1E3D1) and detergent solubilised 1-palmitoyl-2-oleoyl-sn-glycero-3-phosphocholine (POPC, Avanti Lipids) lipid in a 1:10:1000 molar ratio of protein:MSPs:lipids. Following an overnight incubation, the mix was added to activated Biobeads SM2 (400 mg Biobeads: 350 μg of protein) and incubated for 2 hours at 4°C, with rocking, for detergent removal. The nanodisc mix was then transferred with the use of a 25G hypodermic needle (BD Microlance - 0.5×16mm) and loaded onto a His-trap column (HisTrap^TM^ HP, Cytiva) pre-equilibrated with SEC buffer B supplemented with 10 mM imidazole. As a result of the absence of His tag, empty NDs could not bind to the Ni^2+^-resin and were therefore collected in the flow-through (FT), which was collected and then concentrated using a 50 kDa MWCO centrifugal filter. The column, retaining the protein-NDs, was washed with 5 CVs of SEC buffer B containing 30 mM imidazole. Finally, MmpL3-carrying NDs were eluted with 300 mM imidazole buffer and concentrated using a 100 kDa MWCO centrifugal filter. Both samples were then loaded onto a Superdex 200 Increase 10/300 gel filtration column pre-equilibrated with SEC buffer B. Fractions corresponding to main peak were then pooled and concentrated using 100 kDa MWCO centrifugal filters and either used fresh or stored at −80°C for future experiments.

### SDS-PAGE, Native PAGE and Western Blot analysis

For SDS-PAGE analysis, protein fractions were mixed 1: 1 with SDS loading buffer and loaded on a NuPage^TM^ 4-12% Bis-Tris PAGE Gel (Invitrogen™) along with Novex™ Sharp Pre-stained Protein Standards (range from 3.5 to 260 kDa) and separated at a constant voltage of 180 V for 40 minutes in NuPAGE MES SDS running buffer. For native gels, protein fractions were mixed with SDS-free loading buffer in a 3:1 ratio and loaded on a Novex™ 4–20% Tris-Glycine Gels using NativeMark™ Unstained Protein Standards (Invitrogen™) (range from 20 to 1236 kDa) and separated at a constant voltage of 150 V in NativePAGE Running Buffer, for 1 hour or until the dye front reached the bottom of the gel cassette. Denaturing or native gels were either Coomassie-stained and visualised using the GelDoc imaging system by Bio-Rad or rinsed in ultrapure water prior to being used for Western Blot analyses. Native PAGE or SDS PAGE gels were transferred to a nitrocellulose membrane using an iBlot2 (20 V, 7 min; Invitrogen™). The membrane was incubated with blocking buffer (PBS, 0.2% TWEEN 20 and 5% w/v skim milk) for 45 minutes at room temperature, with gentle shaking. Primary anti-His antibody (mouse monoclonal IgG2b, abm^®^) was then added at 1:3000 dilution and membrane was incubated for 1 hour at room temperature. A horseradish peroxidase (HRP)-linked secondary antibody (anti-msIgG2, Cell Signaling Technology) was subsequently added at 1:3000 dilution after extensively washing the membrane with PBS containing 0.2% Tween^®^ 20. After performing three more wash cycles, the signal was developed using Clarity™ Western ECL substrates (BioRad) following the kit manufacturer’s recommendations and the membrane imaged using a GelDoc imaging system.

### Sedimentation velocity analytical ultracentrifugation (SV-AUC)

MmpL3 constructs in SMALPs and NDs samples were prepared to a concentration of 0.1-0.5 mg/mL in 20 mM Tris HCl, 150 mM NaCl, pH 8.0 buffer. Sedimentation velocity experiments were performed at the Birmingham Biophysical Characterisation Facility, University of Birmingham. Experiments were performed at 20°C in a Beckman Coulter XL-I Analytical Ultracentrifuge, using a 50 Ti rotor. Samples were centrifuged for 16 h at 81,000 g, a relative centrifugal force (rcf) that allowed full sedimentation, and monitored by absorbance at 280 nm. SEDNTERP was used to calculate protein partial specific volume, solvent density and viscosity ^22^. Data was then analysed applying a continuous c(s) distribution using the Lamm equation in the program SEDFIT^23^. During the analyses the frictional ratio parameter was allowed to float and the sedimentation coefficient estimates for the particles were normalised and plotted using GUSSI^24^.

### Sucrose gradient separation

Post-SEC concentrated pooled fractions for *Msmeg* MmpL3 ΔC and empty NDs samples in 200 μL of SEC Buffer B (20 mM Tris-HCl, pH 8.0 and 150 mM NaCl) were layered on top of 4 mL linear 10–30% sucrose gradients prepared in the same buffer. Sucrose gradient centrifugations were conducted at 204,000 g in a SW 60 Ti rotor (Beckman Coulter) for 18 h at 4°C with slow acceleration and no brake. Twenty-eight fractions of 150 μL were manually collected and selected aliquots were analysed by Coomassie stained native and SDS–PAGE gels.

### Native mass spectrometry

Prior to analysis by LILBID-MS all analyte samples were buffer-exchanged into 20 mM Tris HCl, pH 8.0 using Zeba Micro Spin desalting columns (Thermo Scientific, USA, 7 kDa MWCO). For each measurement 4 µL of buffer exchanged and degassed protein sample was used. Microdroplets of 50 µm diameter of the analyte were produced by a piezo-driven droplet generator (MD-K-130, Microdrop Technologies GmbH, Norderstedt, Germany) at a frequency of 10 Hz. The microdroplets were transferred to a vacuum and irradiated by a pulsed IR laser operating at the vibrational absorption wavelength of water (2.8 µm). The laser energy, adjusted to 10-23 mJ, is absorbed by the water molecules, resulting in an explosive expansion of the microdroplets and a release of the analyte ions to the gas phase. The analyte ions were accelerated by a pulsed electric field and analysed with a homebuilt time-of-flight mass spectrometer. LILBID settings have previously been published in detail^25,26^. Ion detection was performed in negative mode and data acquisition was carried out using a homebuilt software *Massign* based on LabView^27^. Each mass spectrum shown is the result of an averaged signal of 1000 microdroplets. Data was calibrated based on measurements of an aqueous 10 µM bovine serum albumin solution, smoothed and background corrected.

### Cryo-EM data collection and processing

A 3 μL of aliquot *Msmeg* MmpL3 ΔC NDs at a concentration of 0.6 mg/mL was applied to glow-discharged (Quantifoil Cu R2/2, 300 mesh) holey carbon grids for 30 sec and blotted for 2×5 sec at 4°C and 100% humidity. The grids were plunge-frozen in liquid ethane using a Vitrobot mark IV (Thermo Fisher, Waltham, Massachusetts). Cryo-EM data was collected in the Electron Microscopy Centre at Imperial College London, equipped with a 200 kV Glacios™ 2 Cryo Transmission Electron Microscope (Cryo-TEM) (Thermo Scientific™) with Selectris energy filter and Falcon IV direct electron detector. Cryo-EM images were recorded at −0.7 to 2.5 μm defocus with 2.7 mm spherical aberration and 79 000 x nominal magnification, corresponding to a 1.5 Å/pixel sampling interval. The total specimen dose for each EER fractionated movie was 40 e^−^/A^2^).

The micrographs of MmpL3 were aligned using a patch-based motion correction for beam-induced motion using cryoSPARC^28^. The contrast transfer function (CTF) parameters of the micrographs were determined using Patch CTF^29^. After discarding poor micrographs, with low estimated CTF resolution and thick or crystalline ice, MmpL3 particles were automatically picked using cryoSPARC blob picker and a CHARMM-GUI model of the MmpL3 dimer in 13 nm NDs was used as a template to enrich for dimers. Initially, 1,180,857 particles were selected from 8,946 micrographs after template autopicking in cryoSPARC^28^, using an extract bin x 2 particles. The images were subjected to several rounds of two-dimensional (2D) classification into 200 classes, with an optimised window function to obtain the best protein alignment. False picks and classes with unclear features were removed and resulted in 249,442 particles.

### Lipidomics analysis

BL21(DE3) WT and *ΔacrABEF E. coli* membranes as well as MmpL3 protein in DDM, synthetic and native NDs were subjected to a lipid extraction as described by the Folch method^30^. Internal standards for Cardiolipin (14:0-CDL ammonium salt, #710332), Phosphatidylethanolamine (15:0-18:1-d7-PE, #791638), Phosphatidylcoline (15:0-18:1-d7-PC, #791637), Phosphatidylserine (15:0-18:1-d7-PS, #791639), Phosphatidylglycerol (15:0-18:1-d7-PG, #791640) and Sphingomyelin (18:1-d9 SM, #791649) were purchased from Avanti Polar Lipids, Inc and were added to a final concentration of 10 μg/mL. The lipid extract was separated by liquid chromatography using a Dionex UltiMate3000 RS Autosampler (Thermo Scientific) coupled to a LTQ Velos Pro Orbitrap (Thermo Scientific) mass spectrometer. A 10 μL lipid extract was separated at a flow rate of 0.5 mL/min on an Acuity C18 BEH column (Waters, 50×2.1 mm, 1.7 μm) at 55 °C. Mobile phase A was acetonitrile:water (60:40) with 10 mM ammonium acetate. Mobile phase B was isopropanol:acetonitrile (90:10) with 10 mM ammonium acetate. Mobile phase gradient was linearly changed from 60:40 to 1:99 A:B over 8 minutes and kept constant for 30 seconds before switching back to 60:40 mobile phase A:B over 10 seconds. These conditions were maintained for a further 2.5 minutes. Lipids were analysed by MS in negative ion mode (spray voltage 2.8 kV, desolvation temperature 380 °C, desolvation gas 40 arbitrary units) with range 100-2000 *m/z* and mass resolution 60,000. Lipids were annotated by accurate mass through the LIPID MAPS database^31^. Tandem MS (collision induced dissociation) was performed on membrane samples to fragment intact lipids, in order to identify the fatty acyl chain composition using their diagnostic ions.

### Molecular Dynamics Simulations

*Msmeg* MmpL3 (aa 1-773) oligomeric structures were predicted using the GalaxyHomomer server (http://galaxy.seoklab.org/homomer website)^32^. The algorithm uses templates selected from the protein structure database to predict homomeric structures taking a structure as the input. The MmpL3 dimer model obtained using PDB 7N6B as input and the Resistance-Nodulation-Cell Division (RND) transporter HpnN (PDB ID: 5KHS) as the template was used for further applications ^13,14^. The PDB file for *Msmeg* MmpL3 dimer was processed through the CHARMM (Chemistry at HARvard Molecular Mechanics) - GUI Membrane Builder web-based tool (http://www.charmm-gui.org/)^33^. This tool was used to generate the model of *Msmeg* MmpL3 embedded in a 60% 16:0-18:1 PE (POPE), 20% 16:0-18:1 PG (POPG) and 20% 16:0-18:1 PC (POPC) membrane. PDB was manually oriented with respect to the membrane planes, and the lipid bilayer was then generated by replacement method using the aforementioned ratios of lipid molecules. Furthermore, 0.15 M KCl ions were placed using the Monte Carlo method. All the components were assembled together, including water molecules extending by 22.5 Å above and below the membrane, and a PDB file of *Msmeg* MmpL3 dimer embedded in a lipid bilayer was generated.

AMBER 16 and AmberTools18, including *sander* and the GPU-accelerated *pmemd* code, were used to run MD simulations^34–36^. The PDB file obtained from CHARMM – GUI was initially processed through a Python script (charmmlipid2amber.py) to convert the lipid molecules to be compatible with Amber. All hydrogens were then removed using *pdb4amber*, for the file to be correctly read by the preparatory program *tLeap* ^35^. Force field ff19SB, lipid17 and TIP3P were sourced into *tLeap* to define protein, lipid and water counterparts respectively^37,38^. Finally, topology and coordinate files were generated. MD atomistic simulations of 5 repeats for each model were carried out over a 1.5 μs time frame, after running two minimisations, two heating steps and a 10 ns equilibration. Explicit solvent and 0.15 M KCl were used to solvate the system. Protein was centred within the simulation box and particle mesh Ewald (PME) boundary condition (PBC) applied with a cut off of 10 Å for long-term electrostatic calculations^36^. The two rounds of minimisation were performed using *sander*. For each round, a total of 1000 cycles were carried out starting with the deepest descent algorithm and switching to the conjugate gradient for the last 500 cycles. In the first minimisation step, movements from the protein were restrained using a harmonic potential with a force-constant of 100 kCal mol^−1^ Å^−2^. This restraint was removed for the second minimisation step, allowing the entire system to move freely. Both minimization steps were performed keeping the periodic boundaries volume constant with no pressure scaling and no SHAKE algorithm applied ^39^.

GPU-accelerated *pmemd* code was used to run all the other steps and the SHAKE algorithm was applied to restrain bonds involving hydrogens, therefore allowing the time step used to be increased to 2 fs. The minimised system was heated through two sequential runs to a final temperature of 300 K. First, the system was heated to 100 K over 20 ps, using a Langevin thermostat with random seed and a collision frequency of 1.0 ps^−1^, in a constant volume ensemble (NVE)^40^. During this first heating step a force constant restraint of 10 kCal mol^−1^ Å^−2^ was applied to the protein. The system was then heated to 300 K over 100 ps, by using the same thermostat but this time applying an anisotropic Berendsen weak-coupling barostat to also equilibrate the pressure at 1 bar, thus reproducing a constant T and P ensemble (NPT ensemble). A 10 ns equilibration step followed and was carried out at 1 bar and 300 K. Finally, a 1.5 µs MD production simulation was performed with the same conditions, except for an increased time step of 4 fs, as Hydrogen Mass Repartitioning (HMR) was applied to the solute counterpart^41^.

All trajectories were visualised using the MD movie function in UCSF Chimera. MD trajectories were analysed to identify lipid binding sites on MmpL3. Volumetric maps were generated using the VolMap tool within VMD by calculating headgroup occupancy within 6 Å of the protein over the whole trajectory^42^. Moreover, cpptraj commands *hbond* and *lifetime* were used to investigate protein-protein and protein-lipid hydrogen bonds^43^. Heatmaps were generated using GraphPad Prism 10.0.0 for Mac OS X (GraphPad Software, Boston, Massachusetts USA, www.graphpad.com).

## Results

### Constructs and membrane mimetic systems used in this study

We generated full-length and C-terminal truncated versions of MmpL3 (MmpL3 ΔC) from both *M. smegmatis* and *M. tuberculosis*. The truncated versions of both proteins expressed better than the full-length versions, although we were able to purify all four proteins in DDM and reconstitute them into nanodiscs. We had sufficient pure material for all four proteins in both membrane mimetic systems to allow analysis by SDS-PAGE and Western blot. Native PAGE and SV-AUC were carried out on the *Msmeg* MmpL3 full-length and ΔC in both SMALPs and nanodiscs, while material for *Mtb* MmpL3 ΔC in SMALPs was only sufficient for native PAGE analysis. Only *Msmeg* MmpL3 ΔC in nanodiscs was successfully analysed by LILBID native mass spectrometry and cryo-EM and only this protein was used for MD simulations. Lipidomics analysis was carried out on the *Msmeg* proteins in all three membrane mimetic systems, while only protein in DDM and nanodiscs was analysed for *Mtb* MmpL3.

### Observation of possible oligomers of *M. smegmatis* and *M. tuberculosis* MmpL3 isolated in detergent and following reconstitution into nanodiscs

The first indication that MmpL3 was forming oligomeric arrangements came from SDS-PAGE analysis of full-length *M. smegmatis* MmpL3 (Ms-MmpL3) protein, when isolated in DDM both with/without reconstitution in nanodiscs (see Expanded View Figure 1 for SEC profiles of the isolated material). A band migrating at approximately 110 kDa was observed by both Coomassie blue stain and Western blot analysis (Figure 1a,b, blue boxes) and confirmed as MmpL3 by peptide mass fingerprinting (Mass spectrometry and proteomics facility, University of St Andrews). This band therefore is likely to correspond to the monomeric form of the protein. However, there was a higher molecular weight band (>200 kDa) evident in both the DDM and nanodisc reconstituted samples (Figure 1a,b red boxes). This was also confirmed by peptide mass fingerprinting to be MmpL3. The same pattern of two bands, one low and one high molecular weight was observed for the truncated version of the *M. smegmatis* MmpL3 and both the full-length and truncated versions of the *M. tuberculosis* protein. In our hands we have observed higher molecular weight bands on SDS-PAGE gels that correspond to oligomeric forms of membrane transport proteins^44–47^, as confirmed by structural analysis. However, given the addition of SDS, and the heating that occurs during the electrophoresis process, it was not possible to rule out that these higher molecular weight bands are artefacts of the gel running process.

**Figure 1.**
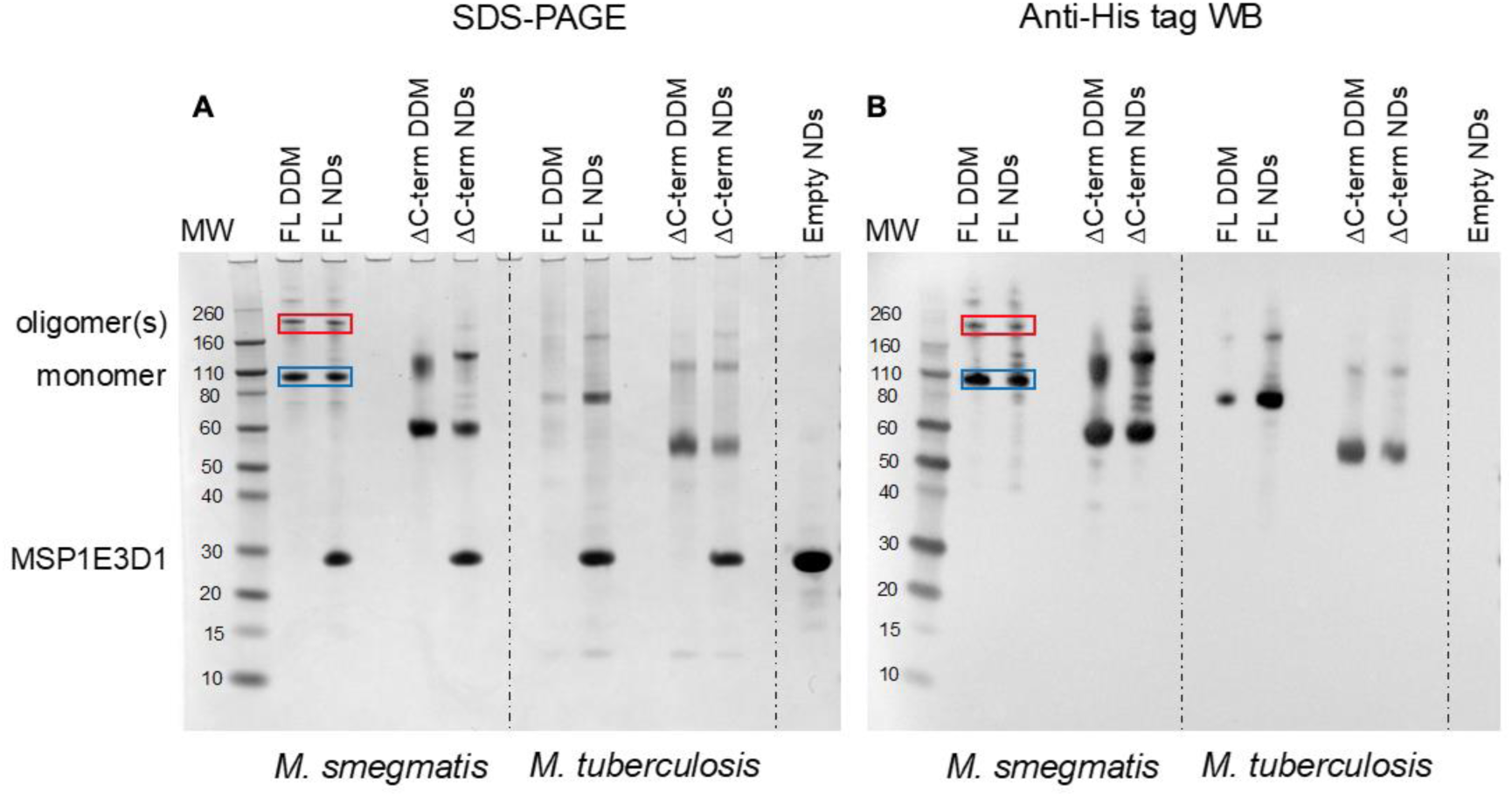
A) SDS-PAGE and B) Western blot analysis of samples obtained during detergent based purification and nanodisc reconstitution of full-length and ΔC MmpL3 from *M. smegmatis* and *M. tuberculosis*. MW = Molecular weight markers, Individual purified proteins in either DDM or nanodiscs (NDs) are indicated above the relevant lanes. The blue and red boxes indicate the likely monomeric and oligomeric forms of the full-length form of MmpL3 from *M. smegmatis* respectively. A similar pattern of two bands is seen for all the other protein constructs.

### MmpL3 is present in two populations in both native and reconstituted nanodiscs

Given that previous structural studies had indicated that the MmpL3 was monomeric, we hypothesised that interactions supporting oligomer formation might be relatively weak and benefit from membrane lipid molecules that are lost during detergent extraction. Thus, we used native nanodiscs for the next step of characterisation of the protein. Native nanodiscs generated by styrene-maleic acid (SMA) polymer extract the protein from the membrane along with an annulus of membrane lipids. There are several examples in the literature where this results in a more physiological arrangement of a protein or protein complex than in detergent based solution^48,49^. Following extraction and isolation of Ms-MmpL3 ΔC in native nanodiscs, SEC analysis revealed the presence of two distinct peaks, one eluting at ∼9-10 mL and one eluting at ∼12 mL (Figure 2A), that were confirmed as two protein populations by native Western blot analysis (Figure 2C). Native Coomassie-stained PAGE analysis and native Western blot analysis (Expanded View Figure 1A) of SEC fractions from both peaks strongly indicated that the first SEC peak (∼9-10 mL, purple bar) contained higher molecular weight protein, migrating between 150 and 250 kDa. The second peak (∼12 mL, red bar) migrating between 60 and 100 kDa is likely to be monomeric MmpL3 protein. Ms-MmpL3 ΔC has a theoretical MW of ∼86 kDa, but migrates at ∼70 kDa on a standard SDS-PAGE gel, as shown in Expanded View Figure 1A. Similar results were obtained for full-length Ms-MmpL3 (Expanded View Figure 1B); the separation of the two SEC peaks was less well defined, but native PAGE confirmed the presence of two distinct populations of protein. The Mtb-MmpL3 ΔC also showed a similar migration pattern of SEC fractions in both SDS- and native PAGE (Expanded View Figure 2).

**Figure 2.**
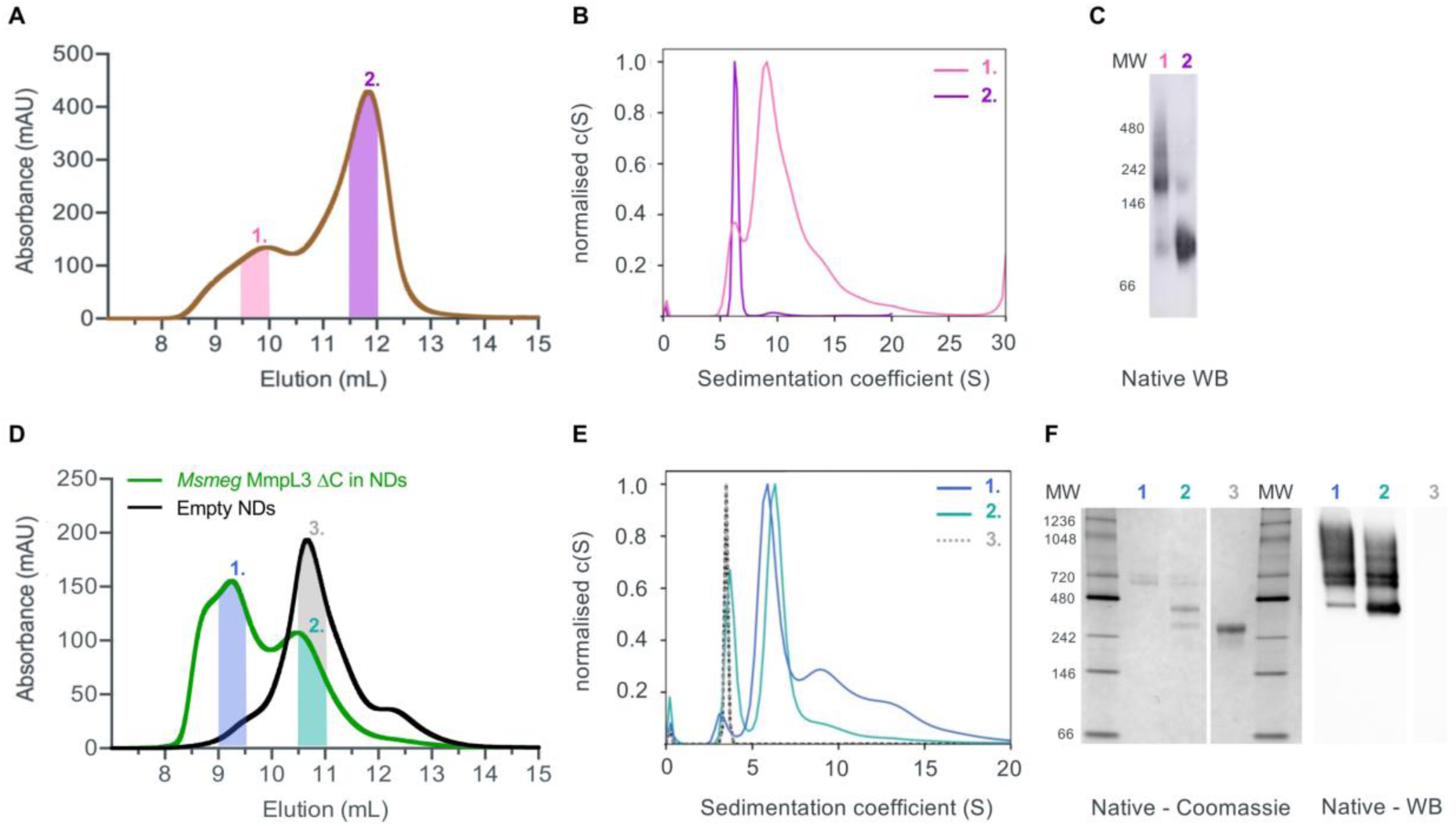
**A** Size exclusion profile of Ms-MmpL3 ΔC isolated in SMALPs. The high (peak 1) and low (peak 2) molecular weight peaks of the MmpL3 protein are indicated by the pink and purple lines. **B** The same colouring is used for the SV-AUC analysis with the traces obtained for protein in peaks 1 and 2 shown in pink and purple respectively. **C** Native Western blot analysis of the SMALP isolated samples using the anti-His tag antibody. **D** Size exclusion profile of Ms-MmpL3 ΔC isolated in DDM and then reconstituted into nanodiscs (green), peaks 1 and 2 are indicated by the blue and cyan lines. The size exclusion profile of empty nanodiscs (black) is shown with the peak (peak 3) indicated by the grey line. **E** SV-AUC profiles of the two populations of Ms-MmpL3 ΔC reconstituted in nanodiscs and empty nanodiscs coloured as in **D**. **F** Native PAGE and native Western blot analysis of the Ms-MmpL3 ΔC reconstituted in nanodiscs and empty nanodiscs. Lanes are labelled with peak numbers.

SV-AUC of the two different populations of the Ms-MmpL3 ΔC confirmed the presence of two species, the predominant species estimated at 126kDa, likely to be monomer and the smaller peak with an estimated MW of 213kDa, possibly a dimer (Figure 2B). It should be noted that SV-AUC of SMALPS does not yield the direct mass of the protein. Instead it provides the mass of the protein alongside the lipid and polymer that makes up the particle. This explains why the masses calculated using SV-AUC are elevated compared to the expected mass. Intriguingly, when we carried out SV-AUC of protein isolated in detergent and then reconstituted into MSP1E3D1 based nanodiscs, we also detected two populations of protein. The difference in size of the two populations was less marked than in the SMALPs, possibly due to the influence of the MSPs on the sedimentation of the samples (Figure 2F).

### Native MS analysis

Both native PAGE and SV-AUC indicated that there are two clear populations of protein; monomer and a likely dimer form of the protein in both SMALPs and nanodiscs. However, these approaches failed to provide definitive proof of the oligomeric status of the protein. For this we used laser induced liquid bead ion desorption (LILBID) MS a soft MS ionisation method, which uses an infra-red laser to drive the release of protein from aqueous based solution allowing the investigation of the stoichiometry of non-covalently bound complexes, among other features, under almost native conditions^26^. The Ms-MmpL3 ΔC in SMALPs proved less stable than that in reconstituted nanodiscs, thus, all further experiments were carried out with

Ms-MmpL3 ΔC in POPC reconstituted nanodiscs. In order to achieve better separation of the different oligomeric forms of the protein than was possible with SEC alone, and to remove as many empty nanodiscs as possible, we also carried out sucrose density gradient (10-30%) ultracentrifugation. As the density increases linearly from top to bottom, different size proteins are fractionated along the gradient, with heavier species and aggregates settling at the bottom of the tube. Complete separation of the different populations in the Ms-MmpL3 ΔC sample was not achieved, as can be seen by the distribution of protein over the 14-24% sucrose gradient range (Figure 3 A, magenta line). However, the empty nanodiscs gave a markedly different separation profile (Figure 3A, black line) compared to the transporter protein sample. Native PAGE and SDS-PAGE analysis (Figure 3B and C) of the Ms-MmpL3 ΔC protein samples obtained at different sucrose concentrations confirmed that empty nanodiscs were present at 12% sucrose, mainly monomeric forms of the protein at 14% sucrose (low sucrose percentage, LSP, sample) and a greater amount of the oligomeric form of the protein observable at 16-18% sucrose (high sucrose percentage, HSP). Both the LSP and HSP samples were subjected to native MS analysis using LILBID and the HSP was used for cryo-EM analysis.

**Figure 3.**
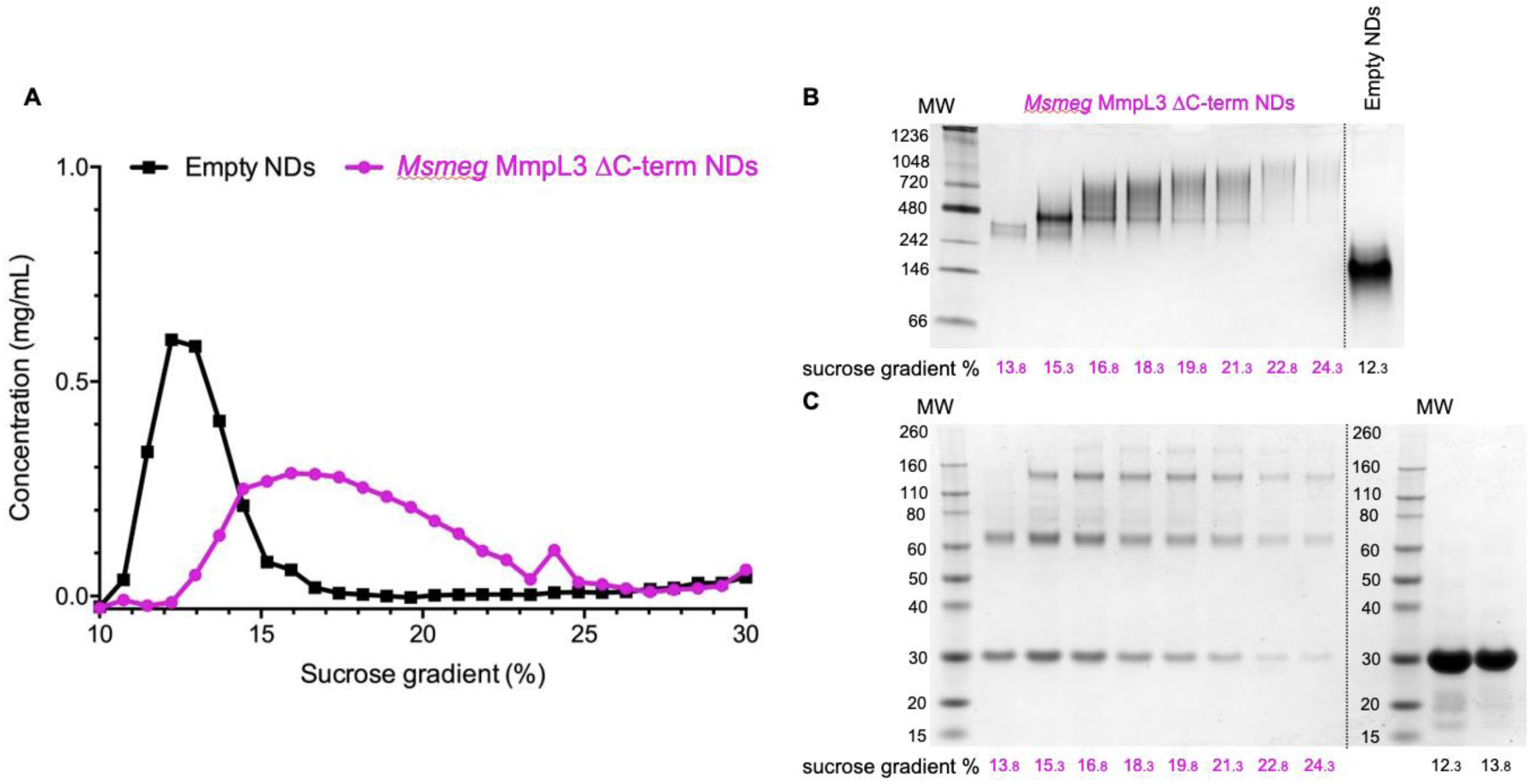
**(A)** Sucrose density gradient ultracentrifugation separation of Ms-MmpL3 ΔC (magenta) and empty (black) nanodiscs over a 10-30% sucrose concentration range. Samples separating at different concentrations of sucrose as indicated were submitted to native PAGE **(B)** and SDS-PAGE **(C)** analysis. The data shown is representative of n = 2 independent experiments.

The LILBID analysis clearly showed that the Ms-MmpL3 ΔC protein that separated to 14% sucrose was monomeric in form with various different charge states. Species with only one MSP have most likely lost the second MSP during the LILBID process. (Figure 4, upper panel). In contrast, the protein sample that migrated into higher concentrations of sucrose (16-18%) was largely dimeric in form again with a variety of charge states (Figure 4, lower panel). This sample also contained small populations of monomeric forms of MmpL3. This may be due to residual monomeric protein in the fraction even after the sucrose gradient separation, or it may derive from oligomers that have dissociated as a result of the native MS procedure. The theoretical masses of complexes including two MSPs and either monomer or dimer of Ms-MmpL3 ΔC are 146,535 kDa and 231,755 kDa, respectively (indicated by dotted lines in the peak insets in Figure 4). The proteo-NDs also contain lipids, some of which are retained during the LILBID process. The broad peaks of the LILBID spectra are due to a statistical distribution of bound lipids in the proteo-NDs. The experimental MWs of the peak insets represent the NDs without any lipids and were determined as 147 kDa and 232 kDa, in very close agreement with the theoretical values and strongly supporting the presence of monomeric and dimeric species in our samples. The dimer was detected at both low and high (Expanded View Figure 3) laser energies.

**Figure 4.**
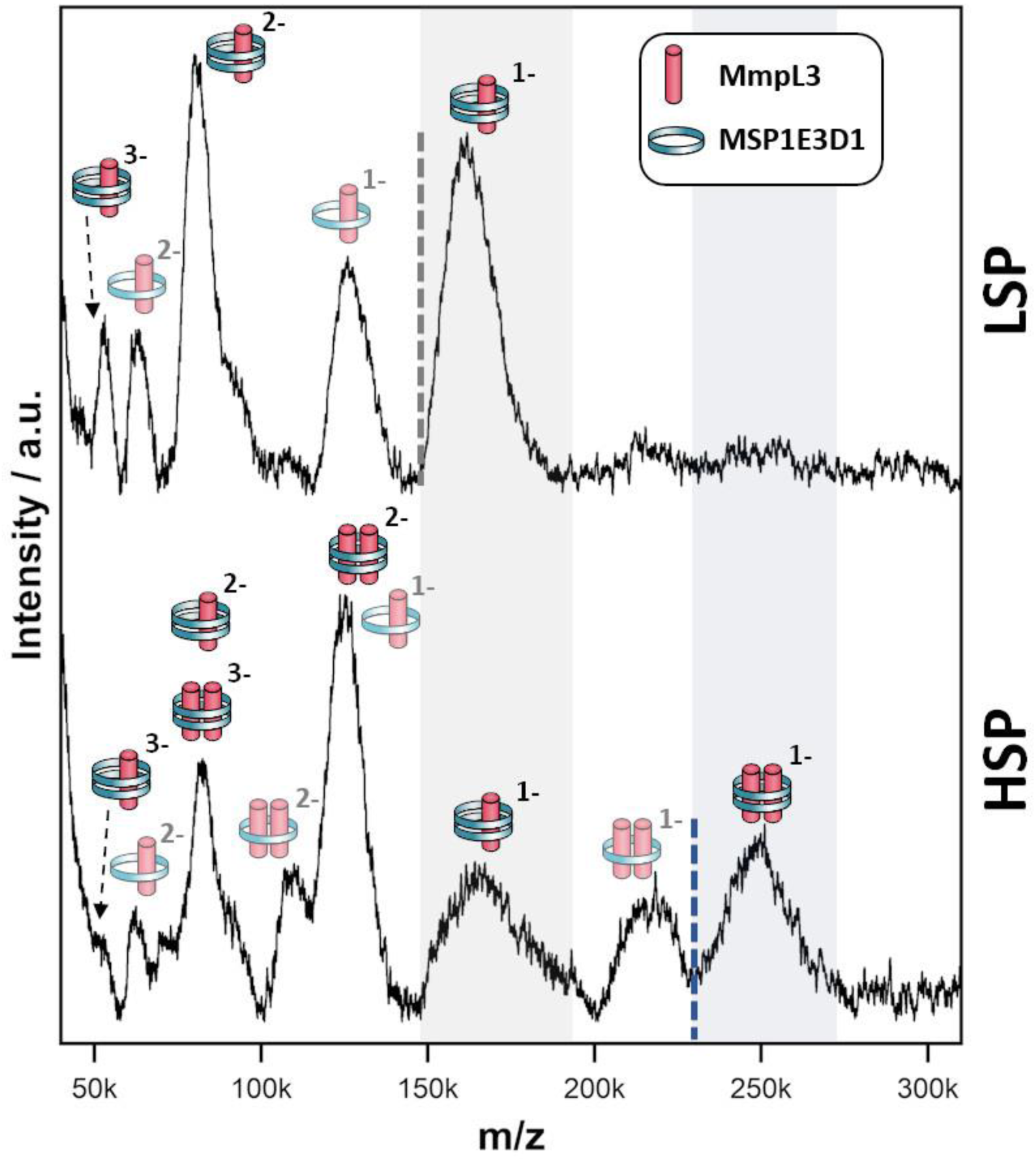
Native Mass Spectrometry for *Msmeg* MmpL3 ΔC NDs. nMS spectra of the *Msmeg* MmpL3 ΔC protein in MSP-NDs separating at low (upper panel) and high (lower panel) sucrose percentage (LSP/HSP) measured at 10 mJ laser intensity. The different protein-MSP combinations detected in each sample and relative charged species are indicated with pictograms. The translucent pictograms are species for which one MSP dissociated during the Laser desorption process. Dotted lines indicate the theoretical mass of NDs particles made of two MSPs copies and either one (grey, panel A) or two (blue, panel B) *Msmeg* MmpL3 ΔC proteins, respectively. The shaded areas to the right of these dotted lines cover the broad peaks, which stem from the inhomogeneous distribution of lipids within the respective proteo-NDs. The grey-shaded area indicates the presence of monomeric MmpL3-NDs while the blue-shaded area indicates dimeric MmpL3-NDs. The data shown is representative of n = 2 independent experiments.

### Cryo electron microscopy

We submitted the HSP sample of Ms-MmpL3 ΔC to cryo electron microscopy and were able to obtain monodispersed samples of both populations of protein (Figure 5A, Extended View Figure 4). In order to facilitate selection of the monomeric versus dimeric protein we generated synthetic models of the monomeric and dimeric forms of the protein in nanodiscs (Figure 5B and C, upper panels) using CHARMM-GUI. Both simulated models show density for the soluble domain on one side of the nanodisc as well as integral membrane protein within the nanodisc, and there is substantially greater density for the dimer. The experimental 2D class averages indicate some likely face on views, which are perpendicular to the dimer interface, with clear differentiation between monomer (Figure 5B, middle panel) and dimer (Figure 5C, middle panel). Views from the side are harder to differentiate since both the monomer and dimer look similar with just slightly greater levels of density when looking through the protein.

**Figure 5.**
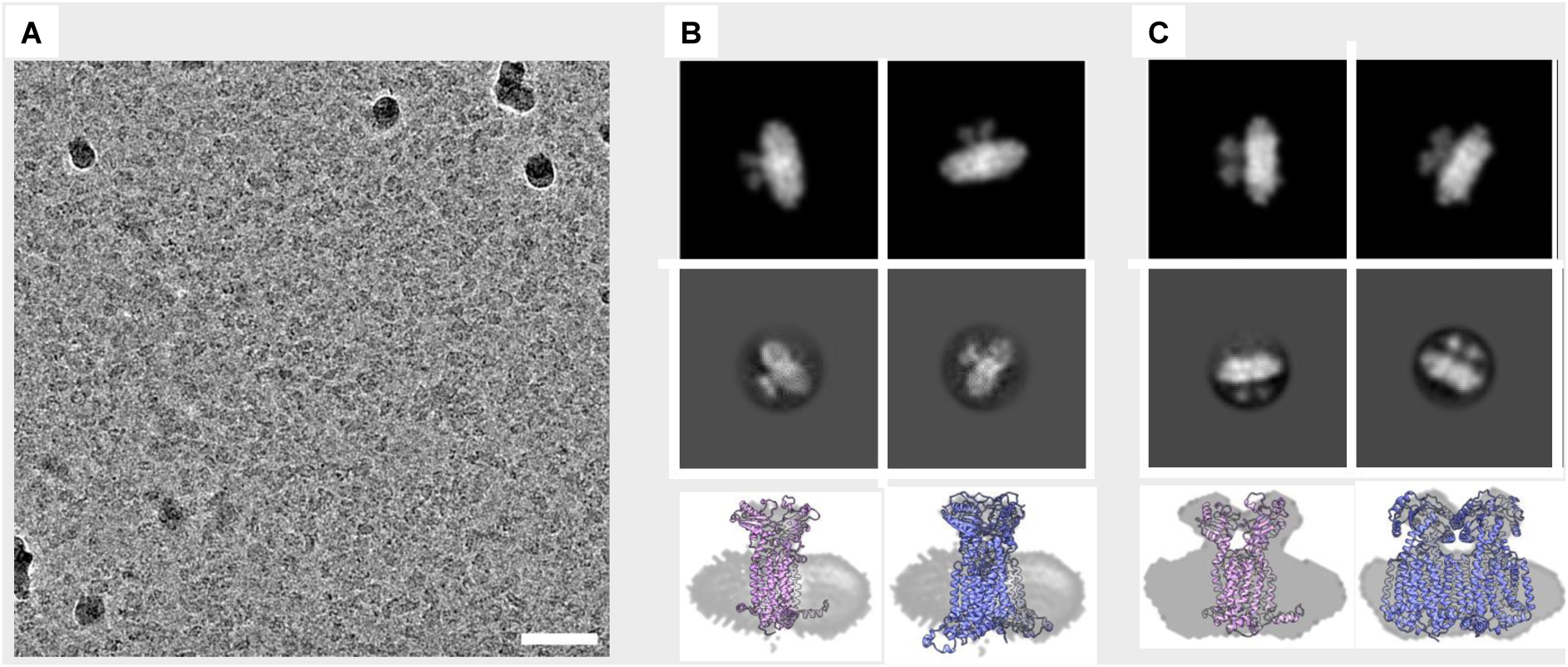
2D classifications of monomeric and dimeric assemblies of MmpL3. Electron cryo micrograph of MmpL3 nanodisc sample, scale bar 200 nm (A). Simulated (top panel) and real (middle panel) 2D classifications of MmpL3 monomer and dimer shown respectively in B and C with bottom panels showing a molecular model fit to side views from 2D classification.

### MD simulations

The resolution from the cryo-EM analysis was too low to provide detailed information on the nature of the interaction and so in order to obtain further insights into the possible organisation of the Ms-MmpL3 ΔC dimer. Experimental analyses as described above were carried out in nanodiscs generated with only POPC. In order to determine a physiologically relevant lipid composition for our MD simulations, we performed lipidomics analysis on MmpL3 constructs. Proteins were isolated in DDM, SMALPs or isolated in DDM and reconstituted into NDs. We identified 59-109 lipids, including from the PE, PC, SM, PG and CDL lipid classes (Extended View Figure 5A). The composition of lipids varied according to the isolation method. Lipids associated with MmpL3 that had been isolated in DDM or SMALPs were mainly from the PE and PG lipid classes. In contrast, when isolated in DDM and reconstituted in NDs, the major lipid class identified was PC. This is expected since NDs are composed of 16:0-18:1 PC lipids (also known as POPC). Next, we calculated the average relative abundance of the main lipid classes across the different constructs/isolation methods (Figure S5B). We therefore chose 60% 16:0-18:1 PE (POPE), 20% 16:0-18:1 PC (POPC) and 20% 16:0-18:1 PG (POPG) as the lipid composition for MD simulations, being representative of the average membrane environment. The MD simulations predict that both the loops connecting specific periplasmic sub-domains (α1-β1, α2-α3, and β2-α4) and TM domain (TM1b-TM2 and TM1a-TM6 helices) regions of the protein are involved in dimer formation (Expanded View Figure 6) and that both Chain A and Chain B contribute H-bond donors and acceptors (Figure 6).

**Figure 6.**
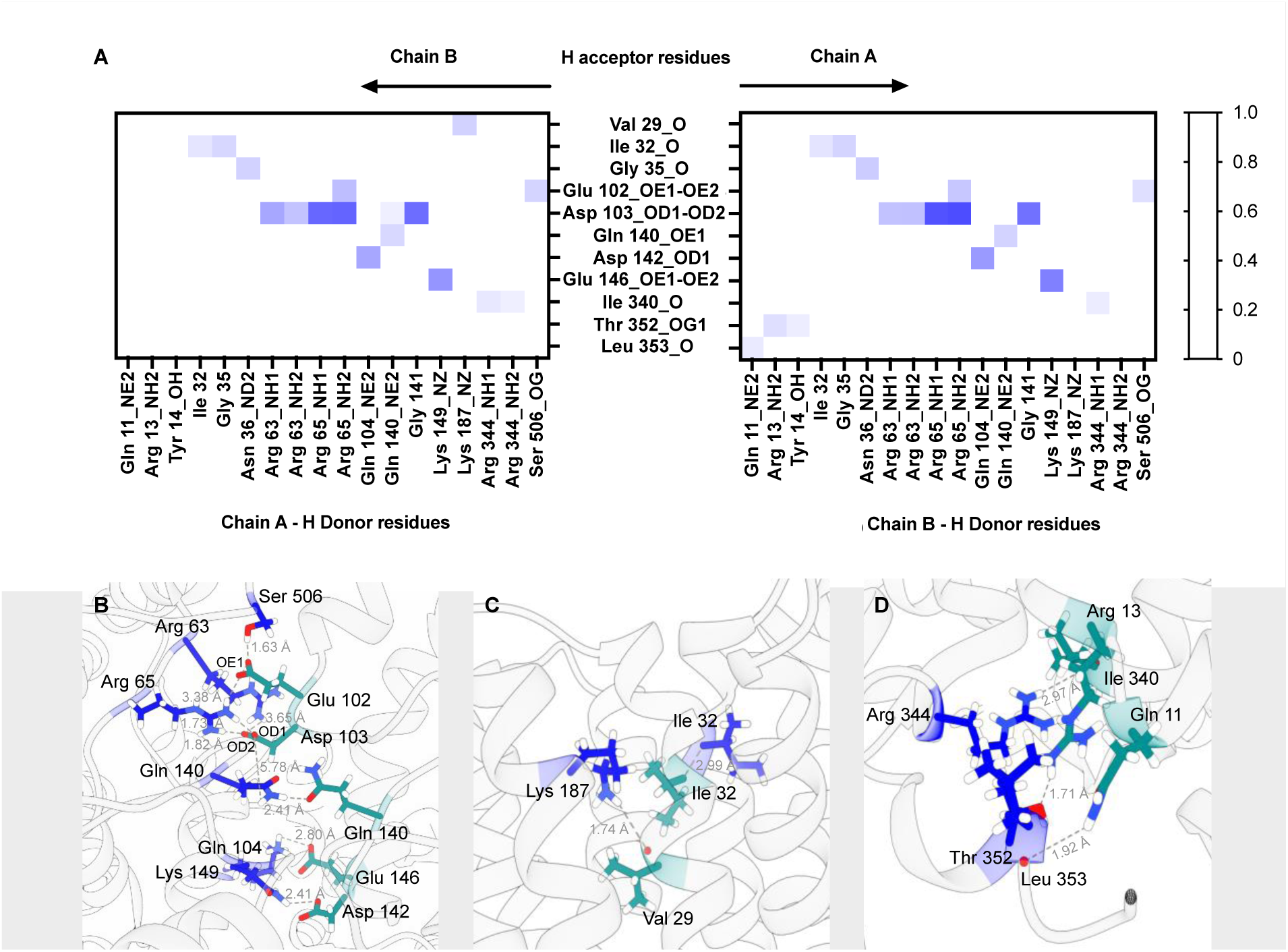
Protein-protein interactions for *Msmeg* MmpL3 dimer. A. Heatmaps for *Msmeg* MmpL3 protein-protein hydrogen bonds using chain A residues as H donors and chain B residues as H acceptors and the reverse. *The* maximum value on the scale shown is 1, indicating that a bond was present in 100% of the simulation frames. The main amino acid residues involved in protein dimerisation are located on periplasmic loops connecting α1-β1 (aa 63-65), α2-α3 (aa 102-104) and β2-α4 (140-149) (B); TM1b (aa 29-32) and TM2 (Lys187) in the region closest to periplasm (C); TM1a (aa 11-13), TM6 (Ile340) and loop connecting TM6 to TM7 (aa 344-353) in region closest to cytoplasm (D). Residues on chain A are in blue while residues on chain B are in dark cyan.

Further MD simulations analysis identified possible interactions between lipid molecules in both leaflets of the simulated membrane, with all three lipid species contributing to the annulus around the MmpL3 dimer (Expanded View Figure 7). It was possible to identify amino acid residues constituting 4 hotspots over the surface of the dimer, two mirrored sites in each protomer (Figure 7A). In our simulation, there are lipids clusters at the lipid substrate entrance date (Figure 7B) comprised of Ser423, Leu424 and His558 of the TM domain and Asn542, Asp531 and Gln554 of the periplasmic domain. We were also able to identify the specific H-bonds formed between MmpL3 and the individual lipid species within the simulated membrane (Figure 7C).

**Figure 7.**
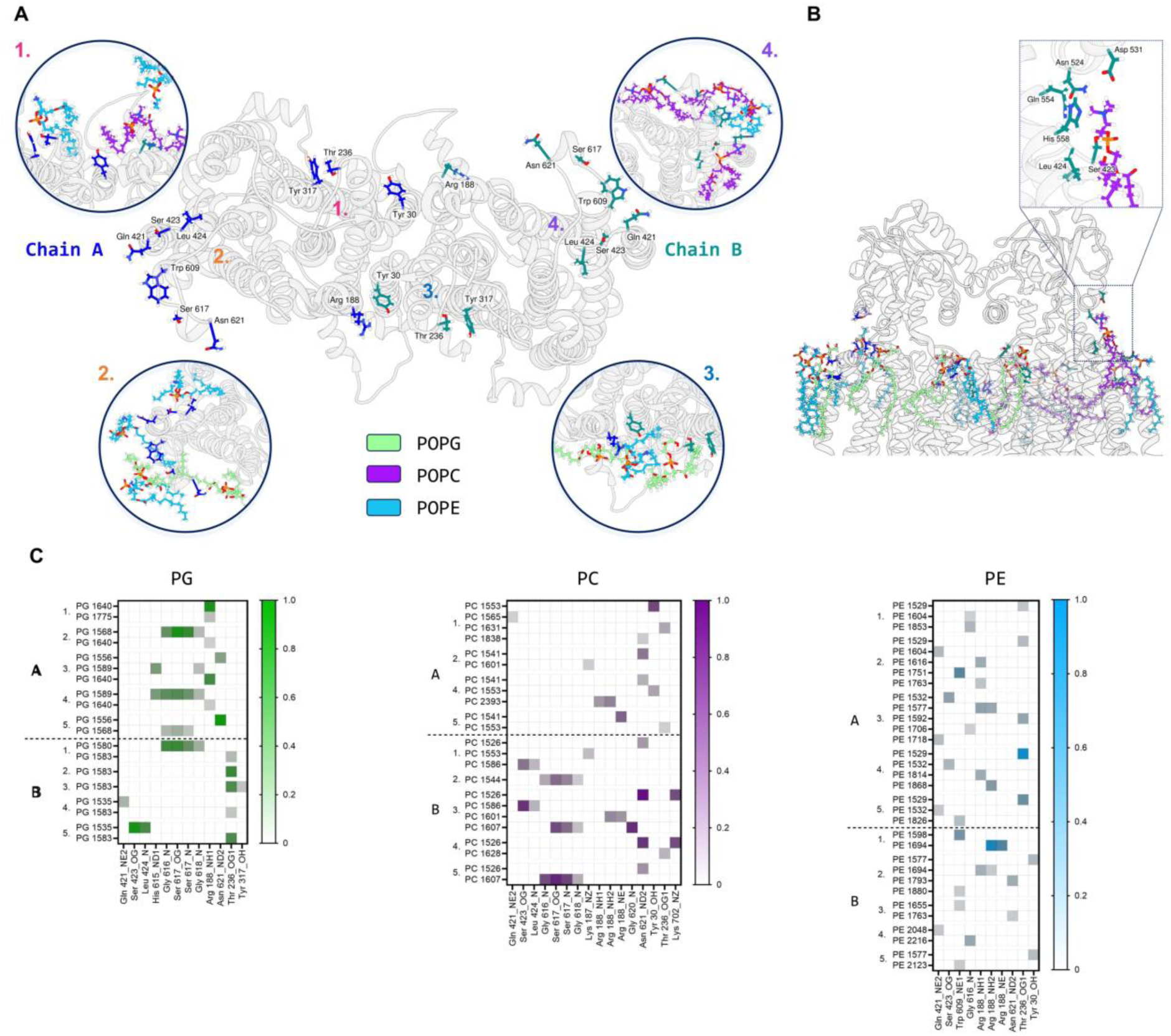
*Msmeg* MmpL3 dimer: protein-lipid interactions on top leaflet. A. Protein top view with main protein residues forming H bonds with lipids labelled and their side chains shown in stick representation. Insets show these residues interacting with lipid species during the simulation (repeat 3 used as representative). B, Protein front view showing lipid species binding to MmpL3 residues, with particular focus on the lipid entrance gate (inset) Relevant residues are indicated. C. Heatmaps for protein-lipids (PG in green, PC in purple and PE in blue) hydrogen bonds by chain and simulation repeat. Protein residues and relative atoms involved in H bond are reported on the x axis, while individual lipid headgroup residues are shown on the y axis. Max value on scale is 1, indicating a 100% of total simulation frames where bond is present. Only interactions >10% of the total frames are plotted.

## Discussion

MmpL3 is an important virulence factor and thus a valid drug target for the development of novel therapies for the treatment of tuberculosis. A full understanding of the structure and mechanism is required for effective drug development. The previous structures of the monomeric forms of the proteins provided insights into the overall structure of MmpL3 protein, the nature of the lipid binding site and the likely mechanism of lipid export ^10–12^.

However, given the well documented oligomeric states of all other RND transporters structurally characterised to date^15,19–21^, it was a little surprising that MmpL3 alone appeared to be monomeric. Early on in our analysis it was evident that MmpL3 could be forming higher oligomeric form states even in DDM-based detergent solution and could be reconstituted into nanodiscs in this form. Our analysis indicates that the reconstituted nanodiscs may stabilise the oligomeric form of the protein. However, the presence of both monomeric and oligomeric states of Mmpl3 was clearest in samples solubilised with SMALPs from the native PAGE and AUC analyses. The increased clarity of the results may bedue to the physiological annulus of lipids that accompanies the protein through extraction and analysis in complex with the SMALPs. However, given that both the monomeric and dimeric forms of the Ms-MmpL3 were most stable in the reconstituted nanodiscs (with POPC), further analysis of the protein was subsequently limited to this form of the protein.

The results from the LILBID and cryo-EM provide very strong evidence that Ms-MmpL3 forms dimers. The limited data we obtained also suggests that this protein adopts the same oligomeric state as the Ms-MmpL3. However, it is important to state that even in our most stringent isolation and separation procedures the dimer was only a proportion of the protein that we were able to obtain. This taken together with the findings from the earlier structural studies indicates that there may be a dynamic equilibrium between the monomeric and dimeric forms.

Our MD simulations indicate a dimer interface that involves interactions between both the TM domain and regions of the periplasmic domain. Polar and charged residues from both regions are involved in the formation of H-bonds between the two protomers. The interaction interface is on the opposite side of the protein to the suggested lipid substrate binding site as indicated by the structure of Ms-MmpL3 in complex with TMM (^11^; PDB: 7N6B). This arrangement both allows the dimer to form and retains access to the binding site as indicated for the related hopanoid transporter, HpnN ^20^. Lipidomics analysis indicates that for all the purification methods, the protein retains a number of different phospholipids. Our MD simulations identified a number of hotspots for lipid binding including the entrance to the substrate translocation channel. Hence it is possible that lipid binding here stabilises a conformation of the protein which ultimately facilitates TMM binding, with the TMM displacing the membrane lipids. Earlier studies have suggested that there is a small but significant rigid body movement of one of the periplasmic domains that is key in substrate transport^11^. In our case, no ligand was present but the periplasmic cavity size reduced at the end of the simulation compared to the beginning with the PD domain of both protomers independently closing.

The possible function of the dimeric arrangement of MmpL3 remains unclear. Typically, the oligomeric states of membrane proteins play roles in function, regulation or stability ^50,51^. The RND transporter AcrB has a trimeric structure with substrate transport coordinated across each of the associated protomers ^17,18^. Structures of the dimer of the hopanoid transporter HpnN from *Burkholderia multivorans* were obtained in two conformational states^20^. These structures highlighted that the large periplasmic domain of the protein undergoes movement suggested to be associated with opening and closing of the substrate translocation channel. Stability required for the movements of this large periplasmic domain may be provided by the dimeric arrangement of the HpnN molecules. Conformational rearrangements of the periplasmic domain associated with TMM translocation through MmpL3 have already been reported^11^, so it is possible that the dimer plays a role in stability in this case too. Our data confirms that by using multi-step purification, reconstitution into nanodiscs and sucrose density gradient ultracentrifugation, a population of dimeric MmpL3 exists that is suitable for high resolution structure analysis.

## Acknowledgements

This research was supported by BBSRC grant BB/V006487/2 awarded to BB. SC was co-funded by an EPSRC PhD scholarship training grant, EP/S023518/1 and the NIHR Imperial Biomedical Research Center.

